# Evaluation of 19 antiviral drugs against SARS-CoV-2 Infection

**DOI:** 10.1101/2020.04.29.067983

**Authors:** Shufeng Liu, Christopher Z. Lien, Prabhuanand Selvaraj, Tony T. Wang

**Affiliations:** Laboratory of Vector-Borne Viral Diseases, Division of Viral Products, Center for Biologics Evaluation, U.S. Food and Drug Administration, Silver Spring, Maryland, United States of America

## Abstract

The global pandemic of the severe acute respiratory syndrome coronavirus 2 (SARS-CoV-2 or 2019-nCoV) has prompted multiple clinical trials to jumpstart search for anti-SARS-CoV-2 therapies from existing drugs, including those with reported *in vitro* efficacies as well as those ones that are not known to inhibit SARS-CoV-2, such as ritonavir/lopinavir and favilavir. Here we report that after screening 19 antiviral drugs that are either in clinical trials or with proposed activity against SARS-CoV-2, remdesivir was the most effective. Chloroquine only effectively protected virus-induced cytopathic effect at around 30 µM with a therapeutic index of 1.5. Our findings also suggest that velpatasvir, ledipasvir, ritonavir, litonavir, lopinavir, favilavir, sofosbuvir, danoprevir, and pocapavir do not have direct antiviral effect.

---

The recently emerged SARS-CoV-2 is a novel coronavirus originated in Wuhan, China (Zhou et al., 2020) and causes coronavirus infectious disease 2019 (COVID-19), a new disease marked by fevers and pneumonia-like symptoms. As of date, SARS-CoV-2 has spread to essentially all continents and killed over 165,000 worldwide. The World Health Organization (WHO) has declared the SARS-CoV-2 outbreak a global pandemic. Unprecedented public-health measures including lockdown of major cities appear to have contained the spread of the virus within China, but the effectiveness of such measures at a global scale remains a contentious topic. Given a prophylactic vaccine will not likely be available in the immediate future, an effective therapeutic intervention could potentially be a game-changer in combating COVID-19. For this reason, multiple countries have unleashed a frenzy of clinical trials hoping to find a drug that can be repurposed to treat COVID-19. Nonetheless, there has been a paucity of information as to the direct antiviral effect of some drugs in trials. Here, we set out an *in vitro* study to evaluate 19 investigational antiviral drugs, including some approved by the FDA for other indications, for their efficacies against SARS-CoV-2. The list includes several viral RNA-dependent RNA polymerase (RdRP) inhibitors, viral protease inhibitors, chloroquine, a furin inhibitor, and a tyrosine kinase inhibitor. The rationale of choosing these drugs was based on *in silico* prediction(Elfiky, 2020), reported possible effect(Liu et al., 2020), and publicly reported trials(Cao et al., 2020). For example, Sofosbuvir, sold under the brand name Sovaldi among others, attracted our attention as a potent, once-daily, orally administered nucleotide analog prodrug inhibitor of the hepatitis C virus (HCV) NS5B polymerase. Sofosbuvir is metabolized *in vivo* to the active antiviral agent GS-461203 (2’-deoxy-2’-α-fluoro-β-C-methyluridine-5’-triphosphate). GS-461203 serves as a defective substrate for the NS5B protein, thus blocks viral RNA synthesis. Sofosbuvir has been identified in molecular docking as a potential SARS-CoV-2 inhibitor(Elfiky, 2020). Favilavir, an approved influenza RdRP inhibitor in China and Japan, is being used in a clinical trial in China to treat COVID-19(Liu et al., 2020). Several HCV or HIV inhibitors, such as Velpatasvir(Chen et al., 2020), Ledipasvir(Chen et al., 2020), Ritonavir(Shah et al., 2020), Lopinavir(Shah et al., 2020) are predicted to be possible anti-SARS-CoV-2 drugs as well.

Our screen assay is based on the protection of SARS-CoV-2-induced cytopathic effect (CPE), which is particularly pronounced in Vero cells. We hypothesized that inhibitors of virus infection would prevent cell death. One major advantage of the CPE-based screen is that compounds that are toxic to cells at the tested doses are simultaneously excluded for future consideration as a drug. To ensure our assay is robust, we carried out the screen under two different infectious doses (multiplicity of infection of 0.01 and 0.001, respectively). For most drugs, we tested three concentrations: 1, 10, 50 µM. When a drug displayed protective effect, we further expanded the range of concentrations to locate the half maximal inhibitory concentration (IC_50_). The results are summarized in Table 1. Briefly, most drugs that we tested failed to prevent SARS-CoV-2-induced cell death, including Sofosbuvir and Favilavir. Additionally, combination of sofosbuvir and velpatasvir or ledipasvir failed to protect infected cells either. Chloroquine, as well as its hydroxyl form, had been allowed by the U.S. FDA to be provided to certain hospitalized patients under an emergency use authorization. In our study, however, chloroquine only showed minimal activity with an IC_50_ close to 30 µM when being added one hour prior to infection with a therapeutic index of 1.5. This finding contrasts with a reported IC_50_ of 6.9 µM in another study(Wang et al., 2020). Given that it normally takes more drug to achieve a full protection in a CPE-based assay, the IC_50_ measured by our assay is expected to be higher than that measured by assays measuring viral RNA(Wang et al., 2020). Nonetheless, when being added post-infection, chloroquine displayed negligible protective effect against virus-induced cell death. Altogether, our finding cautions conducting clinical trials with existing drugs in treating COVID-19. Without even *in vitro* efficacy support, using these drugs in patients is risky because potential side effects. Indeed, ritonavir and lopinavir were declared ineffective after a failed clinical trial(Cao et al., 2020); Brazilian government recently halted a small trial involving chloroquine after patients developed irregular heart rates. Our results did support the potential of Remdesivir as a drug to treat COVID-19. Remdesivir (GS-5734) is a nucleotide prodrug that has broad antiviral activity against viruses from different families in vitro(Lo et al., 2017), and therapeutic efficacy in nonhuman primate models of lethal Ebola virus and Nipah virus infection(Lo et al., 2019; Warren et al., 2016). Remdesivir (GS-5734) effectively inhibited MERS coronavirus (MERS-CoV) replication in vitro and showed efficacy against Severe Acute Respiratory Syndrome (SARS)-CoV in a mouse model(Sheahan et al., 2017). In consistent, compassionate use of remdesivir in patients hospitalized for severe COVID-19 displayed clinical improvement in 68% of the patients (68%)(J. Grein, 2020). Overall, our study reinforces the notion that clinical trials must be supported by rigorous pre-clinical studies as we take an accelerated pace of developing new drugs to combat SARS-CoV-2.

**Table 1.** Evaluation of 19 investigational antiviral drugs against SARS-CoV-2 Infection.

| Drug Name | Known antiviral mechanism | Multiplicity of Infection |  | IC<sub>50</sub> (μM) | CC<sub>50</sub> (μM) | SI | Concentrations tested (μM) |
| --- | --- | --- | --- | --- | --- | --- | --- |
|  |  | 0.01 | 0.001 |  |  |  |  |
| *Chloroquine pretreatment* | antimalarials and amebicides | + | + | 30 | 45 | 1.5 | 0.25,0.5,1,2,5,10,20,30,60,125,250,500 |
| *Chloroquine posttreatment* | antimalarials and amebicides | ± | ± | >50 | 45 | <0.9 | 0.25,0.5,1,2,5,10,20,30,60,125,250,500 |
| *Remdesivir* | nucleotide prodrug | + | + | 2.5 | 175 | 70 | 0.025, 0.05, 0.1, 0.2, 0.4, 0.8,1.5, 3.125, 6.25, 12.5,25, 50,100, 200 |
| *Sofosbuvir* | HCV RdRP inhibitor | - | - | - | >200 | - | 0.01,0.1,0.5,1,2,5,10,50,100,150,200 |
| *Velpatasvir* | HCV NS5A inhibitor | - | - | - | 17 | - | 0.025, 0.05, 0.1, 0.2, 0.4, 0.8,1.6, 3.125, 6.25, 12.5,25, 50 |
| *Ledipasvir* | HCV NS5A inhibitor | - | - | - | - | - | 1, 10, 50 |
| *Sofosbuvir+Velpatasvir* | HCV RdRP inhibitor + NS5A inhibitor | - | - | - | - | - | 1, 10, 50 |
| *Sofosbuvir+Ledipasvir* | HCV RdRP inhibitor + NS5A inhibitor | - | - | - | - | - | 1, 10, 50 |
| *Danoprevir* | HCV protease inhibitor | - | - | - | >50 | - | 0.025, 0.05, 0.09, 0.195, 0.39, 0.78,1.56, 3.125, 6.25, 12.5,25, 50 |
| *Nelfinavir* | HIV protease inhibitor | - | - | - | - | - | 1, 10, 50 |
| *Ritonavir* | HIV protease inhibitor | - | - | - | - | - | 1, 10, 50 |
| *Saquinavir* | HIV protease inhibitor | - | - | - | - | - | 1, 10, 50 |
| *Indinavir* | HIV protease inhibitor | - | - | - | - | - | 1, 10, 50 |
| *Amprenavir* | HIV protease inhibitor | - | - | - | - | - | 1, 10, 50 |
| *Lopinavir* | HIV protease inhibitor | - | - | - | - | - | 1, 10, 50 |
| *Raltegravir* | HIV integrase inhibitor | - | - | - | - | - | 1, 10, 50 |
| *Tenofovir* | nucleotide analog reverse-transcriptase inhibitor | - | - | - | - | - | 1, 10, 50 |
| *AZT* | HIVs reverse transcriptase inhibitor | - | - | - | - | - | 1, 10, 50 |
| *Favilavir* | influenza RdRP inhibitor | - | - | - | - | - | 1, 10, 50 |
| *Pocapavir* | Poliovirus capsid inhibitor | - | - | - | 12.5 | - | 0.025, 0.05, 0.1, 0.2, 0.4, 0.8,1.6, 3.125, 6.25, 12.5,25, 50 |
| *Bemcentinib (R428)* | tyrosine kinase AXL inhibitor | - | ± | - | - | - | 1, 10, 50 |
| *Chloromethylketone* | Furin convertase inhibitor | - | - | - | - | - | 1, 10, 50 |

-, 0-10% protection; ±, 10-40% protection; +, >50% protection under at least one concentration that was tested. TI, therapeutic index (CC<sub>50</sub>/ IC<sub>50</sub>).

## Methods

### Drugs, Cells and Virus

Remdesivir, sofosbuvir, danoprevir, pocapavir, bemcentinib (R428) were purchased from Medchemexpress LLC. Chloroquine, velpatasvir, ledipasvir were bought from Fisher. Nelfinavir, ritonavir, saquinavir, indinavir, amprenavir, lopinavir, raltegravir, tenofovir, AZT were from NIH AIDS Reagent Program. Favilavir was obtained from Cayman Chemical and chloromethylketone was from Enzo Life Sciences.

Vero E6 cell line was purchased from American Type and Cell Collection (ATCC) and cultured in eagle’s minimal essential medium (MEM) supplemented with 10% fetal bovine serum (Invitrogen) and 1% penicillin/streptomycin and L-glutamine.

The SARS-CoV-2 strain WA1-2020 was obtained from BEI Resources, NIAID, NIH, and had been passed three times on Vero cells and 1 time on Vero E6 cells prior to acquisition. It was further passed once on Vero E6 cells in our lab.

### Virus titration

SARS-CoV-2 strain WA1-2020 was titered using the Reed & Muench Tissue Culture Infectious Dose 50 Assay (TCID50/ml) system (Reed, 1938). Vero cells were plated the day before infection into 96 well plates at 1.5 × 10^4^ cells/well. On the day of the experiment, serial dilutions of virus were made in media and a total of 6-8 wells were infected with each serial dilution of the virus. After 48 hours incubation, cells were fixed in 4% paraformaldehyde followed by staining with 0.1% crystal violet. The TCID50 was then calculated using the formula: log (TCID50)=log(do)+ log (R)(f+1). Where do represents the dilution giving a positive well, f is a number derived from the number of positive wells calculated by a moving average, and R is the dilution factor.

### CPE-based drug screening assay

VeroE6 cells were seeded at 1.8 ×10^4^cells/well in a 96 well plate 24 hours before the experiment. After cells had reached 80% confluency, they were infected with SARS-CoV-2 in the presence or absence of a compound for two hours. One hour (or whatever time it is) after the infection the virus was removed, and fresh compound was added back to each well. Two Days after infection media was removed from the cells and at least 50μl of the CellTiter-Glo® Reagent was then quickly added to all wells in the luminometer plate and the luciferase reading was quantified with a Veritas luminometer (Promega). The cytopathic effect was quantified by the CellTiter-Glo Luminescent Cell Viability Assay system (Promega, WI). Both IC_50_ and CC_50_ were calculated using Prism 7.0 (Graphpad).

## Acknowledgements

We thank Dr. Wilson, Mr. LaClair and Ms. Howard for their support throughout the study. We are grateful to Drs. Weir and Feigelstock for editing. The following reagent was deposited by the Centers for Diseases Control and Prevention and obtained through BEI Resources, NIAID, NIH: SARS-Related Coronavirus 2, Isolate USA-WA1/2020, NR-52281.

The authors declare no conflict of interest.

